# Cellular determinants of metabolite concentration ranges

**DOI:** 10.1101/439232

**Authors:** Jeanne M. O. Eloundou-Mbebi, Anika Küken, Georg Basler, Zoran Nikoloski

## Abstract

Cellular functions are shaped by reaction networks whose dynamics are determined by the concentrations of underlying components. However, cellular mechanisms ensuring that a component’s concentration resides in a given range remain elusive. We present network properties which suffice to identify components whose concentration ranges can be efficiently computed in mass-action metabolic networks. We show that the derived ranges are in excellent agreement with simulations from a detailed kinetic metabolic model of *Escherichia coli*. We demonstrate that the approach can be used with genome-scale metabolic models to arrive at predictions concordant with measurements from *Escherichia coli* under different growth scenarios. By application to 14 genome-scale metabolic models from diverse species, our approach specifies the cellular determinants of concentration ranges that can be effectively employed to make predictions for a variety of biotechnological and medical applications.

**Author Summary:** We present a computational approach for inferring concentration ranges from genome-scale metabolic models. The approach specifies a determinant and molecular mechanism underling facile control of concentration ranges for components in large-scale cellular networks. Most importantly, the predictions about concentration ranges do not require knowledge of kinetic parameters (which are difficult to specify at a genome scale), provided measurements of concentrations in a reference state. The approach assumes that reaction rates follow the mass action law used in the derivations of other types of kinetics. We apply the approach with large-scale kinetic and stoichiometric metabolic models of organisms from different kingdoms of life to show that we can identify a proportion of metabolites to which our approach is applicable. By challenging the predictions of concentration ranges in the genome-scale metabolic network of *E. coli* with real-world data sets, we further demonstrate the prediction power and limitations of the approach.

## Introduction

Advances in systems biology studies have been propelled by the availability of high-quality genome-scale metabolic reconstructions for many organisms across all kingdoms of life [1]. Metabolic network reconstructions contain information about metabolites and reactions through which they are transformed to support different cellular processes [2, 3]. Alongside enzyme concentrations and phenomenological constants, reaction rates and metabolite concentrations—as two faces of the metabolic phenotype—characterize key aspects of the metabolic capabilities of an organism. Since metabolic concentrations are important determinants of reaction rates [4], understanding what controls their physiological ranges can point to cellular mechanisms that control phenotypic plasticity to ensure viability of organisms under changing conditions [5].

A naïve approach to determine a concentration range for a given component is to assume that it is present with a single molecule or that the entire cell dry weight under an investigated scenario is composed solely of that component. This derivation requires only knowledge of the component’s molecular weight, which is readily available. However, the derived ranges are vast and largely invariant under different scenarios; therefore, they are not informative. Here we ask whether large-scale metabolic models can be used for accurate prediction of concentration ranges. Resolving this question is tantamount to identifying a cellular mechanism underlying the control of concentration range for given cellular component.

The change in concentration of metabolites can be described by a system of coupled ordinary differential equations (ODEs), 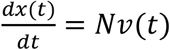, where *v*(*t*) = (*v*_1_(*t*),…, *v*_*n*_(*t*))^*T*^ denotes reaction rates and *x*(*t*) = (*x*_1_(*t*),…, *x*_*m*_(*t*)) the metabolite concentrations at time *t*, and *N* represents the stoichiometric matrix. The rows of the stoichiometric matrix correspond to metabolites, columns stand for reactions, and its entries denote the stoichiometric coefficients with which metabolites participate in reactions as substrates or products [6]. Reaction rates are modeled according to a kinetic law, *v*(*t*) = *f*(*x*(*t*), *θ*), which often leads to nonlinearities and involves multiple parameters, denoted by *θ* [7]. As a result, the coupled nonlinear ODEs are often not analytically tractable and their simulations are challenging. These issues arise since parameters remain poorly specified at a genome scale for the majority of model organisms [8, 9] and the nonlinear ODEs may lead to numerical issues [10]. In addition, determining the steady-state concentration ranges by characterizing the solutions to the system of non-linear equations *Nf*(*x*(*t*), *θ*) = 0 is intractable for large-scale networks even when the equations have a simplified mass action form often used in metabolic modeling [11].

Feasible steady-state reaction rates, *v*, for which *Nv* = 0, can be predicted based solely on the structure of the network with computational approaches from the constraint-based modeling framework [12]. However, since intracellular reaction rates cannot be measured directly, the validation of these predictions requires laborious labeling experiments and model fitting procedures [13]. By neglecting the effect of concentrations on reaction rates, constraint-based approaches do not facilitate the usage of metabolic network reconstructions to predict concentrations of metabolites, which are becoming more accessible by quantitative metabolomics technologies [14].

The existing constraint-based approaches that can make predictions of metabolite concentrations and their ranges are based on consideration of thermodynamics constraints. Thermodynamics-based metabolic flux analysis (TMFA) produces flux distributions that do not contain any thermodynamically infeasible reactions or pathways, and it provides information about the free energy change of reactions and the range of metabolite concentrations in addition to reaction fluxes [15]. However, due to uncertainty in the estimation of the standard Gibbs free energies, TMFA usually predicts unconstrained ranges for metabolite concentrations (see Discussion in Henry et al. [15]). Metabolic Tug-of-War (mTOW) extends TMFA but is based on a non-convex optimization approach which comes at a cost of local optima and lack of robustness of predictions (validated by correlation [16]). A method to predict metabolite concentration ranges with limited knowledge about the underlying kinetic laws and parameter values would allow direct integration and validation of genome-scale models with experimental data from metabolomics technologies, enabling systems biology applications, from engineering of intervention strategies to design of new drugs [17–19].

Here we provide an approach which relies on the structure of the network, encoded in the stoichiometric matrix, to provide simulation-free prediction of steady-state concentration ranges by employing mass action kinetics. We focus on mass action kinetics since it underlies the derivation of more involved types of kinetics for different reaction mechanisms [20], allows for consideration of enzyme concentration if enzymes appear as reaction substrates, and provides a simple mathematical form that may be amenable to analytical treatment. The usage of mass action was here also favored due to lack of information about reaction mechanisms at a genome-scale level. The approach expands on the well-established concept of full coupling of reactions [21] to consider pairs of reactions whose ratio of mass-action-compatible fluxes depends only on the respective rate constants. Thus, this flux ratio is invariant at any, not necessarily steady, state of the system. The approach is also refined to predict concentration ranges for unseen cellular scenarios provided concentration data from a reference experiment. Our method complements the constraint-based modeling framework, focused on analysis of steady-state reaction rates, and thus enables a comprehensive characterization of feasible concentrations in genome-scale metabolic networks under specified conditions.

## Results

### Metabolites with structurally constrained concentrations (SCC)

Consider a metabolic network composed of *m* metabolites that participate in *n* reactions. The (*m* × *n*) stoichiometric matrix, *N*, can be written as a difference of two non-negative matrices, *N* = *N*^+^ − *N*^−^, where *N*^+^ includes the stoichiometry of the products and *N*^−^ comprises the stoichiometry of the substrates of each reaction. For instance, the stoichiometry of substrates and products given in Fig. 1b describes the metabolic network on Fig. 1a. We assume that the rate of reaction *R*_*i*_ is modeled according to mass action kinetic, whereby 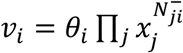, where *θ*_*i*_ > 0 is the reaction constant and the concentration of each substrate molecule appears in *v*_*i*_ as a multiplicative factor.

**Fig. 1.**
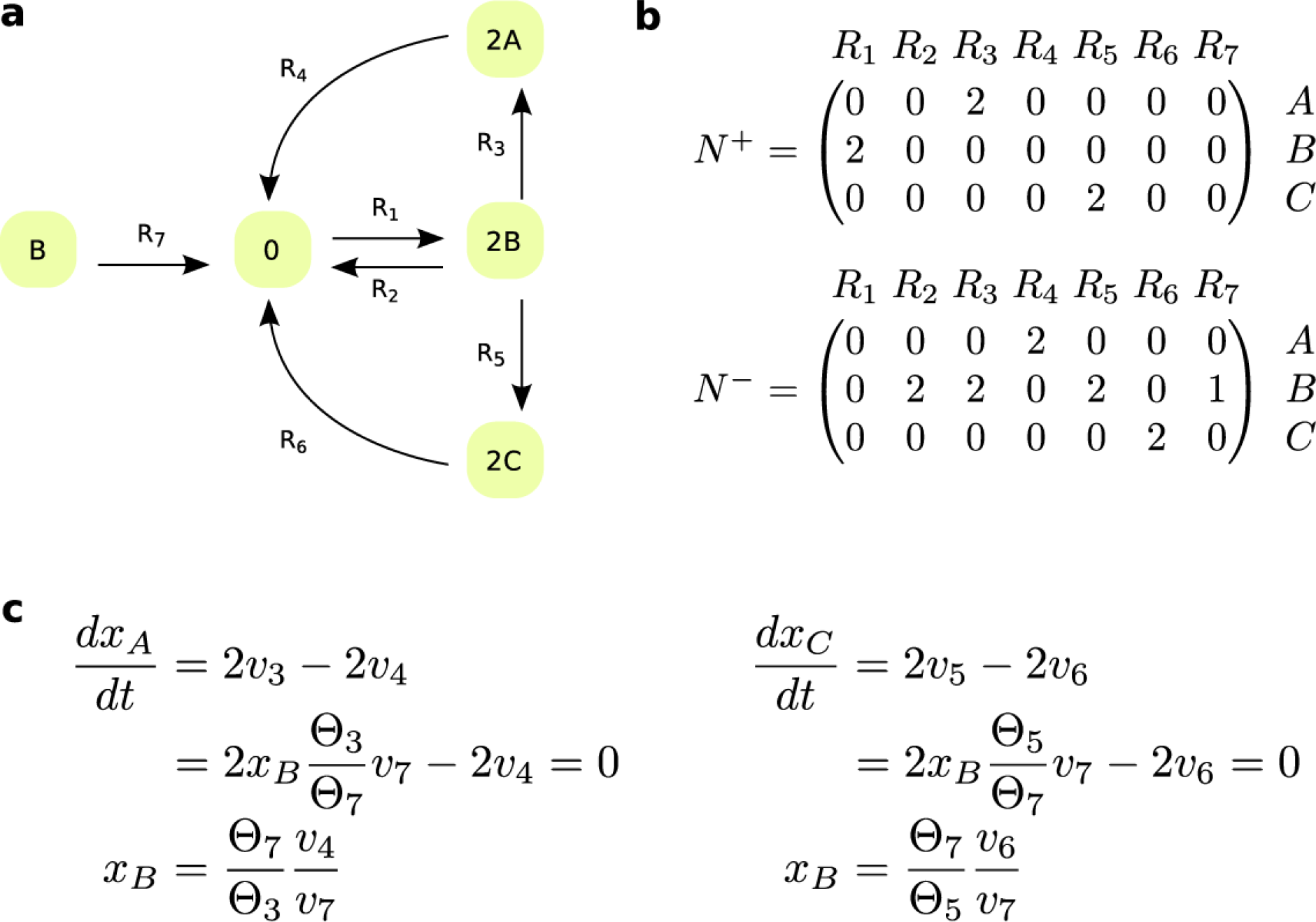
Network with a component exhibiting structurally constrained concentration. (**a**) Reaction diagram that includes seven reactions, *R*_1_ - *R*_7_, and three components, *A* - *C*. (**b**) stoichiometric matrices associated with substrates, *N*^−^, and products, *N*^+^, for the network in (a). Reaction *R*_7_ lacks one substrate molecule of *B* in comparison to *R*_2_, since 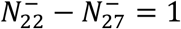 and 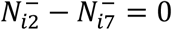for every *i* ≠ 2. Reactions *R*_3_ and *R*_5_ share the same substrate components with same stoichiometry, and hence their fluxes are fully coupled under the assumption of mass action kinetic. Reaction *R*_3_ is fully coupled to reaction *R*_4_, as are reactions *R*_5_ and *R*_6_. (**c**) Component *B* exhibits structurally constrained concentration from the ODEs of components *A* and *C*. The network exhibits different positive steady states with changing rate of reaction *R*_1_.

To state our main result, we require some concepts and terminology which we next introduce and illustrate. We will say that a reaction *R*_*k*_ lacks one substrate molecule of *X*_*i*_ in comparison to reaction *R*_*l*_, if 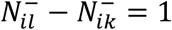 and for every 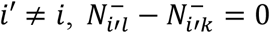. For the network in Fig. 1a, reaction *R*_7_ lacks one substrate molecule of component *B* in comparison to reaction *R*_2_. Under the assumption of mass action kinetic, if a reaction lacks one substrate molecule in comparison to another, the reactions differ in their orders by one. As a result, the ratio of fluxes for such reactions at any state of the system depends only on the rate constants and the concentration of the substrate in which the reactions differ.

Furthermore, two reactions *R*_*k*_ and *R*_*l*_ are fully coupled if there exists *λ* > 0, such that *v*_*l*_ = *λv*_*k*_ for any positive steady-state reaction rate *v*, *i.e.*, *Nv* = 0 [21]. Therefore, fully coupled reactions have an invariant ratio over all positive steady states that the network admits, and full coupling is a transitive relation. For the network in Fig. 1a, reaction *R*_3_ is fully coupled to *R*_4_ and *R*_5_ is fully coupled to *R*_6_. Such reactions, which are fully coupled irrespective of the kinetic law, can be efficiently determined based on the stoichiometry of large-scale networks by linear programming [21, 22] (see Materials and Methods).

Under the assumption of mass action kinetic, two reactions that share the same substrates of same stoichiometry are also fully coupled [23]. In this case, the coupling holds for any, not necessarily steady, state of the system. Therefore, the consideration of mass action kinetic expands the set of fully coupled reactions. For instance, this is the case for reactions *R*_3_ and *R*_5_ that have the substrate components of same stoichiometry in Fig. 1a, whereby 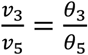.

Consider now a metabolite *X*_*j*_ with an ODE given by 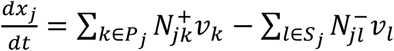 where *P*_*j*_ is the set of reactions with *X*_*j*_ as one of their products and *S*_*j*_ is the set of reactions which have metabolite *X*_*j*_ as one of their substrates. A metabolite *X*_*i*_, not necessarily different from *X*_*j*_, has structurally constrained concentration (SCC), if the following conditions hold: (*i*) for each reaction *R*_*l*_ in *S*_*j*_, there exists a non-empty subset 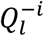 of reactions lacking one substrate molecule of *X*_*i*_ in comparison to *R*_*l*_; the union of all 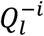 yields the set of reactions 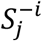; (*ii*) all reactions in 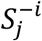 are mutually fully coupled; and (*iii*) all reactions in *P*_*j*_ are mutually fully coupled. A similar argument can be made with respect to condition (*i*) in terms of reactions in the set *P*_*j*_ (Materials and Methods). A metabolite *X*_*i*_ that satisfies the conditions above will be referred to as a SCC metabolite.

In the following, we use the ODE for metabolite *X*_*j*_ to derive the concentration bounds for a metabolite *X*_*i*_ with SCC. Let *Q* be a subset of 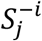 that contains one and only one reaction from each of 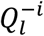. Under mass action, for the flux of every reaction *R*_*l*_ ∈ *S*_*j*_, it then holds that 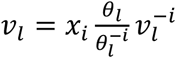 (see Materials and Methods), where 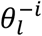 is the reaction constant and 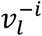 the flux of a reaction 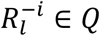. The expression for 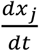 above then becomes 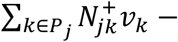 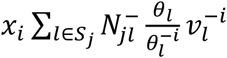.

At any positive steady state, it then holds that 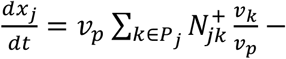 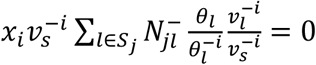, for any flux *v*_*p*_ of reaction *R*_*p*_∈ *p*_*j*_ and flux 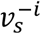 of reaction 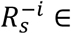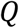. Due to the conditions (*iii*), above the sum 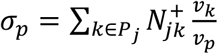 is a constant which in the simplest case, when all reactions in *P*_*j*_ are fully coupled irrespective of the kinetic rate law, depends only on the network structure. In addition, due to condition (*ii*), above, the value of 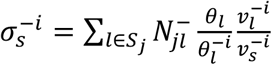 is also a constant which depends on both the network structure and a subset of rate constants. The rate constants which appear in the expression for 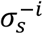 and *σ*_*p*_ for any 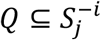 will be referred to as *relevant rate constants*, while the flux ratio 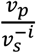 will be called *relevant flux ratio*.

Therefore, given a steady-state flux distribution, *v*, a set 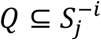, and two reactions *R*_*P*_ ∈ *P*_*j*_ and 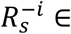, we have that 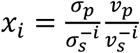. This derivation establishes a direct relation between a flux distribution, under specified inputs from the environment, and the concentration of a SCC metabolite. We can also use the derived expression to obtain the concentration bounds for *x*_*i*_ over any set, *F*, of steady-state flux distributions and subset *Q* (per definition above), yielding the following: 
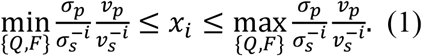

For instance, component *B* in Fig. 1a is SCC, derived from the ODE of component *A*, whereby the relevant flux ratio is 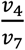 and the relevant rate constants are *θ*_3_ and *θ*_7_ (Fig. 1c). Similarly, one can show that component *B* is SCC from the ODE of component *C*.

Let the lower and upper bounds for the concentration of metabolite *X*_*i*_ derived from the ODE of metabolite *X*_*j*_ in Eq. (1) be denoted by 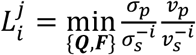 and 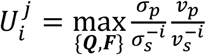 respectively. If there are *r* metabolites *X*_*d*_, 1 ≤ *d* ≤ *r* for which Eq. (1) applies, then the lower and upper bounds for the concentration of *X*_*i*_ are given by the intersection of the ranges derived from the ODEs of *X*_*d*_, i.e‥ 
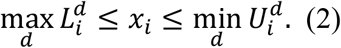

Therefore, the lower bound is the minimum of the maxima, while the upper bound is the maximum of the minima derived from the individual ODEs. In case that the SCC of a metabolite can be derived from multiple ODEs, Eq. (2) provides more constrained predictions about metabolite concentration ranges than Eq. (1) alone. For instance, component *B* in Fig. 1a is SCC not only from the ODE of component *A* but also from that of *C*, whereby the relevant flux ratio is 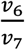 and the relevant rate constants are *θ*_5_ and *θ*_7_ (Fig. 1c). In case that the upper bound is smaller than the lower bound in Eq. (2) then the system of ODEs does not have a positive solution for *X*_*i*_, which implies that the network does not allow a positive steady state. Therefore, the approach can also be used to check for existence of positive steady state with respect to a SCC metabolite under mass action kinetics.

### Validation of the approach with a large-scale kinetic model of *E. coli*

The proposed approach can be employed to determine metabolite concentration ranges by using information about full coupling of reactions, fluxes entering relevant flux ratios, and the relevant reaction rate constants. To validate the predictions, we employ a detailed kinetic model of elementary metabolic reactions of *E. coli* [8] from which these inputs are readily available. Of the 830 metabolites interconverted by 1,474 elementary reactions in the model, our approach determines that 23 metabolites exhibit SCC. The ranges for these SCC metabolites are fully determined by 67 relevant rate constants (4.6% of all rate constants) and fluxes of 67 reactions (4.6% of all reactions) which enter in the relevant flux ratios. We use the kinetic model to simulate 100 steady states from different initial conditions (Supplementary Table S1).

We determined the Euclidean distance between the predicted and simulated lower and upper bounds to demonstrate their quantitative agreement. Since metabolite concentrations vary over several orders of magnitude, the results based on Euclidean distance will be biased by the presence of very large metabolic pools; therefore, we also considered two variants of relative Euclidean distance (see Materials and Methods). Our results from the quantitative comparison demonstrate a very good agreement between the predicted and simulated bounds (Supplementary Table S2, Supplementary Fig. S1). We also employ the Pearson correlation to assess if the predicted and simulated bounds agree qualitatively across the metabolites with SCC. We determine that there is a perfect qualitative match between the predicted and simulated lower (1, p-value < 10^−6^) and upper bounds (1, p-value < 10^−6^) of the SCC metabolites (Supplementary Table S2).

It has been recently proposed that the shadow prices of metabolites can be used to quantify the ranges of metabolite concentrations, under the assumption that the cellular system optimizes an objective [24]. To compare the performance of shadow prices as a measure of metabolite concentration ranges, we employ the stoichiometric matrix of the analyzed kinetic model by using the maximization of metabolic exchange fluxes as cellular objective, shown to outperform yield as a predictor of growth rate [25]. We did not use optimization of yield, most widely used in flux balance analysis, since the model has been parameterized without consideration of a biomass reaction. We observe that for the analyzed model and the physiologically relevant objective, the calculated shadow prices for the 23 SCC metabolites cannot be used as indicators of concentration variability due to the weak negative correlation with the concentration ranges as well as with the coefficients of variation of the SCC metabolites (Supplementary Table S2). These findings point out that our approach, in absence of a cellular objective but with knowledge about a few rate constants and selected flux ratios, outperforms the existing contender for quantifying concentration ranges in large-scale metabolic networks.

### Effects of missing information about rate constants

While the full reaction couplings considered by our approach can be readily obtained given the structure of the network and flux ratios are increasingly available from labeling approaches [26], the resulting predictions can be affected by missing information about rate constants. To assess the effect of missing information on the accuracy of predictions, we consider the cases that 10 - 90% of rate constants used in the derivation of the ranges for the metabolites with SCC are known (see Methods). We consider three scenarios whereby the missing ratios of rate constants, appearing in Eq. (1), are substituted by: (*i*) a value of one, simulating a scenario in which all relevant rate constants are of the same value, (*ii*) the mean, or (*iii*) the median of the ratios of relevant rate constants that are present (*i.e.*, known) in the model equation from which the conditions for SCC are established. We note that the units of the rate constants are not relevant since rate constants enter Eq. (1), above, as ratios.

We find that the substitutions for the missing ratios of rate constants according to the three scenarios, as expected, decrease the Pearson correlation between predicted and simulated ranges over 100 instances of models in which relevant rate constants were removed at random (Fig. 2). Nevertheless, even when only 30% of the relevant rate constants are known for the cases (*i*) and (*iii*), we obtain a median Pearson correlation coefficient between the predicted and simulated ranges of at least 0.6 (Fig. 2). Substituting the missing ratio of rate constants with the mean of the ratios shows the largest variability over the 100 instances of models with partial knowledge of rate constants. The reason for this finding is that the distribution of rate constants and their ratios are highly left-skewed (Supplementary Fig. S2). Therefore, we conclude that even in the absence of information about rate constants that matches the current state-of-the-art of knowledge about *E. coli* (Supplementary Table S3), our approach provides qualitatively reliable estimates of concentration ranges in large-scale models. The ordering of lower and upper bounds between metabolites can be predicted well (median significant Spearman correlation above 0.75 at significance level of 0.05 for all scenarios). However, we observe that the median over relative and log-transformed Euclidean distances between predicted and simulated lower as well as upper bounds over the 23 SCC metabolites are small (<0.71 and <0.08, respectively) when more than 50% of the relevant rate constants are known (Supplementary Fig. S3-S6). Therefore, the approach can be used for the frequently employed comparison of metabolite concentration ranges within and between conditions.

**Fig. 2.**
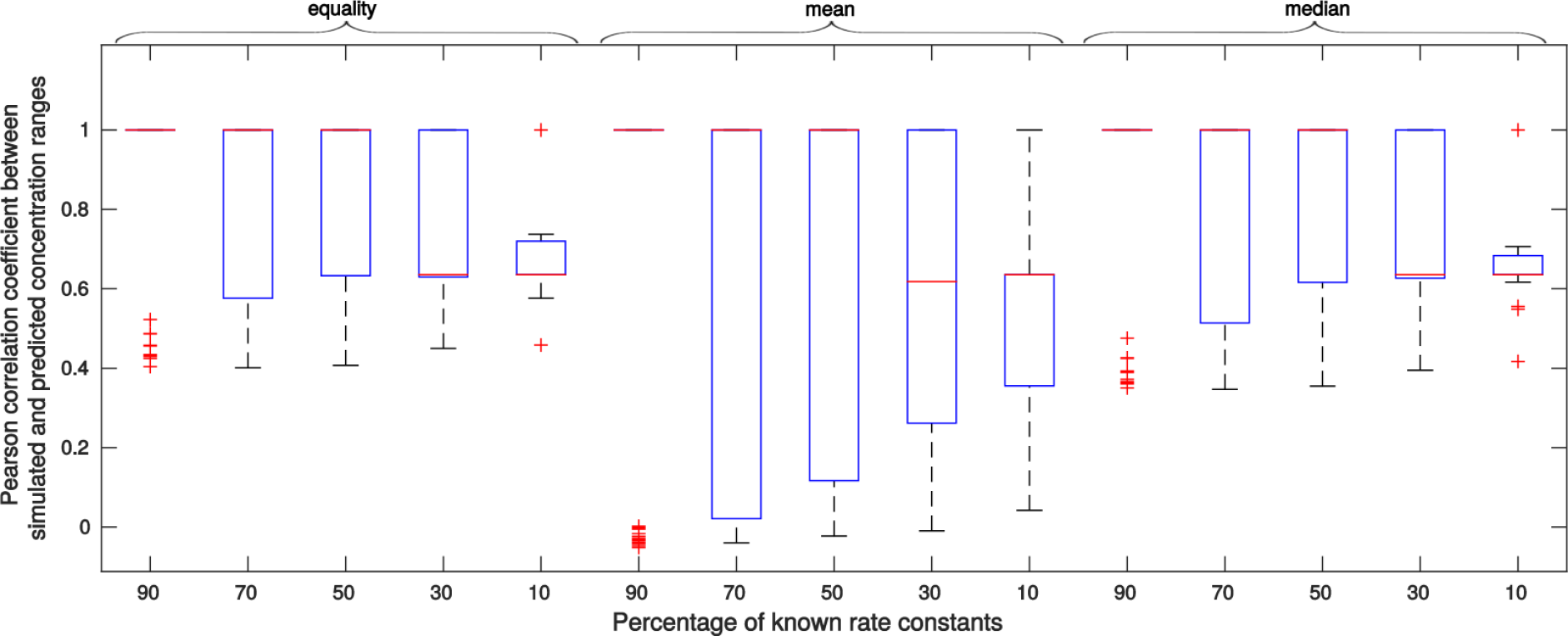
Effect of missing information about relevant rate constants on the accuracy of concentration range predictions for a large-scale kinetic model of *E. coli*. We consider 10-90% of the relevant rate constants to be unknown by random removal. We consider three scenarios for the substitution of missing ratios of rate constants: (*i*) equality (i.e., kinetic rate constants are assumed to be the same), (*ii*) the mean, or (*iii*) the median of the ratios of relevant rate constants that are still present in the model. Shown are the boxplots (red lines inside each box denote the corresponding medians) of the resulting Pearson correlation coefficients between the predicted and simulated ranges over the SCC metabolites in the kinetic model of *E. coli*.

### Effect of missing information about flux ratios

We also investigate the accuracy of the predictions of concentration ranges when full information about relevant rate constants is available and relevant flux ratios are obtained from constraint-based modeling approaches. To obtain physiologically relevant predictions, we constrain the model with the simulated exchange fluxes (Supplementary Table S1), since they can be readily obtained from experiments (e.g. by following substrate depletion). As the employed kinetic model does not specify a biomass reaction, we optimize a weighted average of ATP production and total flux, known to lead to predictions in line with flux estimates from labeling experiments [2]. To this end, we determine the range for the relevant flux ratios at the optimal value for the objective and used them together with Eq. (2) to obtain concentration ranges for the 23 SCC metabolites (Materials and Methods). We find that for 13 out of 23 SCC metabolites the predicted concentration range reside inside the respective simulated range. For additional 6 metabolites the ranges overlap, while the remaining metabolites show no overlap in the predicted and simulated range using the objective of optimized ATP production and total flux (Fig. 3). Since the approach provided accurate quantitative and qualitative predictions with perfect information in the case of kinetic modeling, the discrepancy is due to the objective used to constrain the physiologically reasonable fluxes.

**Fig. 3.**
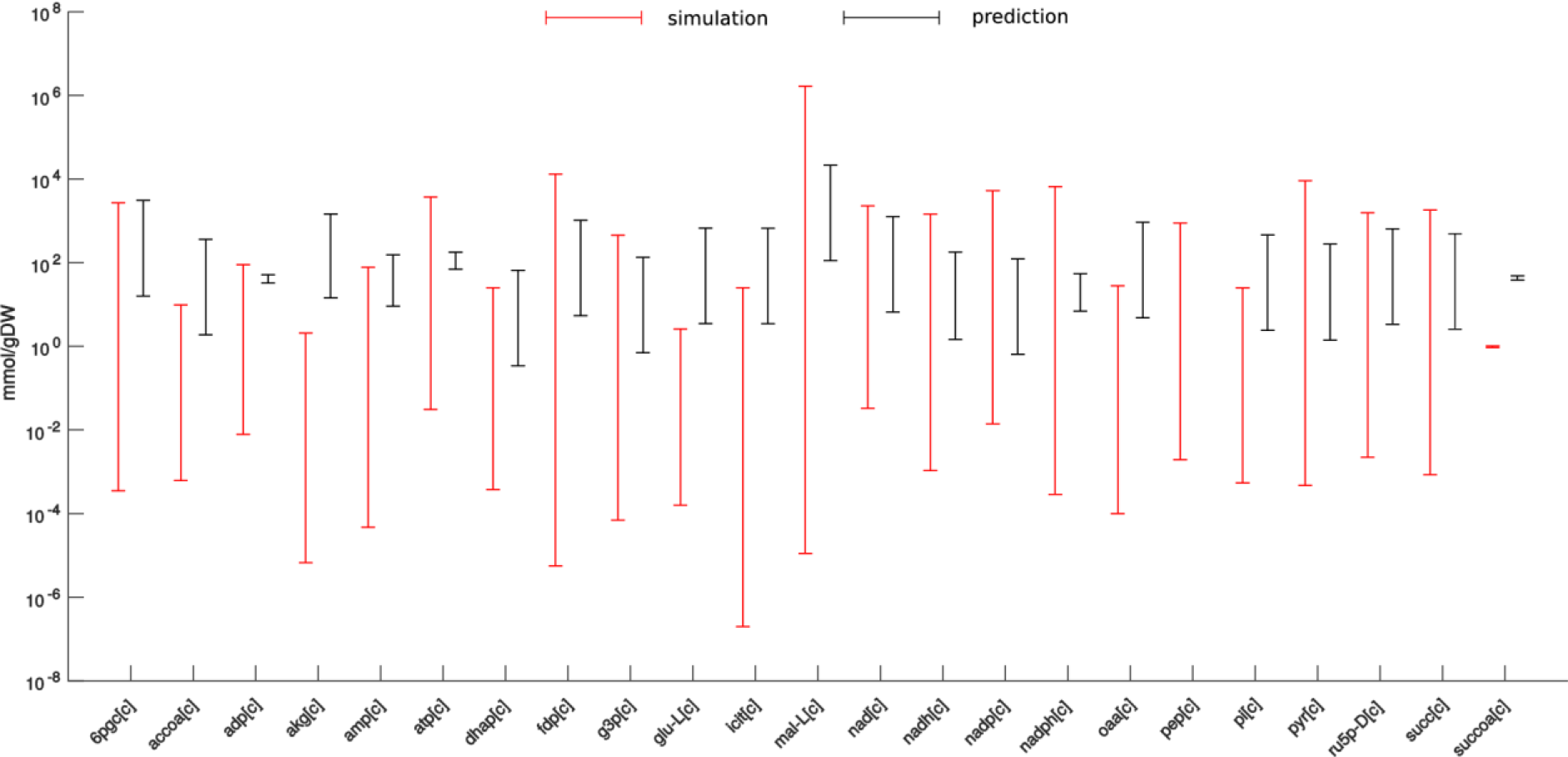
Effect of missing information about relevant flux ratios on the accuracy of concentration range predictions for a large-scale kinetic model of *E. coli*. Relevant flux ratios are obtained by constraint-based modeling in which the objective of weighted ATP production and total flux is maximized. Red bars denote simulated ranges resulting from 100 different initial conditions of the large-scale kinetic model of *E. coli*. Black bars denote the predicted ranges following Eq. (2). Concentration ranges are predicted for 23 SCC metabolites in the employed metabolic model.

### Concentration ranges in a genome-scale metabolic model of *E. coli*

Arguably the most interesting scenario for application of our approach is with genome-scale metabolic networks. We find 199 SCC metabolites in the cytosol and 168 in the periplasm and extracellular space of the most recent genome-scale metabolic network of *E. coli* [27] (Supplementary Table S8). However, for this model, we observe that there are data available for only 28% of relevant rate constants (Supplementary Table S3), and we have no estimates of the relevant flux ratios available from labeling experiments [28–30]. Therefore, the approach cannot be used without extensions. Given a steady-state flux distribution, *v*, the concentration of a SCC metabolite *X*_*i*_ is given by 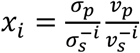. If we have data on concentration of SCC metabolites and flux predictions from the constraint-based modeling framework, we can readily obtain estimates for the ratio 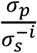. By definition, this ratio is invariant over the conditions where all steady-state fluxes appearing in relevant flux ratios are non-zero. Therefore, we can use the estimates for 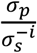 together with flux predictions to make predictions about concentration ranges following Eq. (2) for another scenario. We note that the prediction about concentration ranges inherit the uncertainty in the estimation of 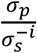 as well as the flux ratios from flux balance analysis, which may contribute to the size of the predicted ranges.

#### Metabolite concentration data set of Ishii et al. [28]

We use the measurements of steady-state concentrations of 182 metabolites from *E. coli* under different growth scenarios [28]. This data set includes 15 of the 199 cytosolic SCC metabolites found in the genome-scale model. We also have access to rates of glucose and oxygen uptakes, carbon dioxide release as well as growth from the same experiments [28], which we use as constraints to a genome-scale metabolic network of *E. coli*. It has been shown that *E. coli* does not optimize a single objective (e.g., growth), but its steady-state flux distributions result from the trade-off between tasks of optimizing growth, ATP synthesis, and total flux [2]. Since growth rate is fixed from measurements, we optimize the weighted average of ATP synthesis and total flux, with a weighting factor of 0.1 on ATP synthesis to reduce the effect of the order difference in the respective optimum observed when ATP production and total flux are optimized individually. Here, too, at the obtained optimum we can efficiently estimate ranges for the relevant flux ratios (Materials and Methods). In addition, we compare obtained concentration ranges with those predicted when maximization of ATP is used as the only objective. To obtain estimates for 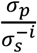, we use three replicates for the concentration data and predictions of ranges for relevant flux ratios at growth rate of 0.2*h*^−1^ (Supplementary Table S4). Eq. (2) can then be applied to determine concentration ranges based on 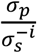 for a combination of replicates, to investigate the effect of outliers. We predict in turn the concentration ranges for three other growth rates (i.e., 0.4, 0.5, and 0.7*h*^−1^).

For the objective of optimizing ATP synthesis and total flux, our results demonstrate that measurements for 9, 10, and 6 of the 15 SCC metabolites fall in the predicted concentration range for the three growth rates, respectively (Fig. 4). Nevertheless, the Spearman correlation between the measured values and the predicted lower and upper bounds is significant and larger than 0.57 and 0.56, respectively (Supplementary Table S5). Therefore, the approach can be used to compare the ordering of lower or upper bounds between different experimental scenarios (Supplementary Fig. S7). In addition, this analysis highlights the effect of the replicates of metabolite concentrations used in calculating the values of 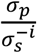, since estimates for some of the replicates may be outliers (Fig. 4). In contrast, we find that 4, 5 and 2 of the 15 SCC metabolites fall in the measured range for the three growth rates when maximization of ATP is used as objective (Supplementary Fig. S9). Moreover, we cannot predict concentrations for 8 out of the 15 SCC metabolites due to numerical instabilities arising when using this objective under the additionally imposed constraints on growth. The reasons for the discrepancy between the predicted and measured values under both objectives include the combination of at least three factors: the inability to distinguish the concentrations of free metabolites from those bound to macromolecules experimentally [31], model (and objective) inaccuracies, and the simplifying assumption of mass action kinetic. Nevertheless, the approach can be extended to consider networks with kinetic laws derived from mass action which involve enzyme forms (e.g., Michaelis-Menten, see Discussion) at cost of increased data requirements for application.

**Fig. 4.**
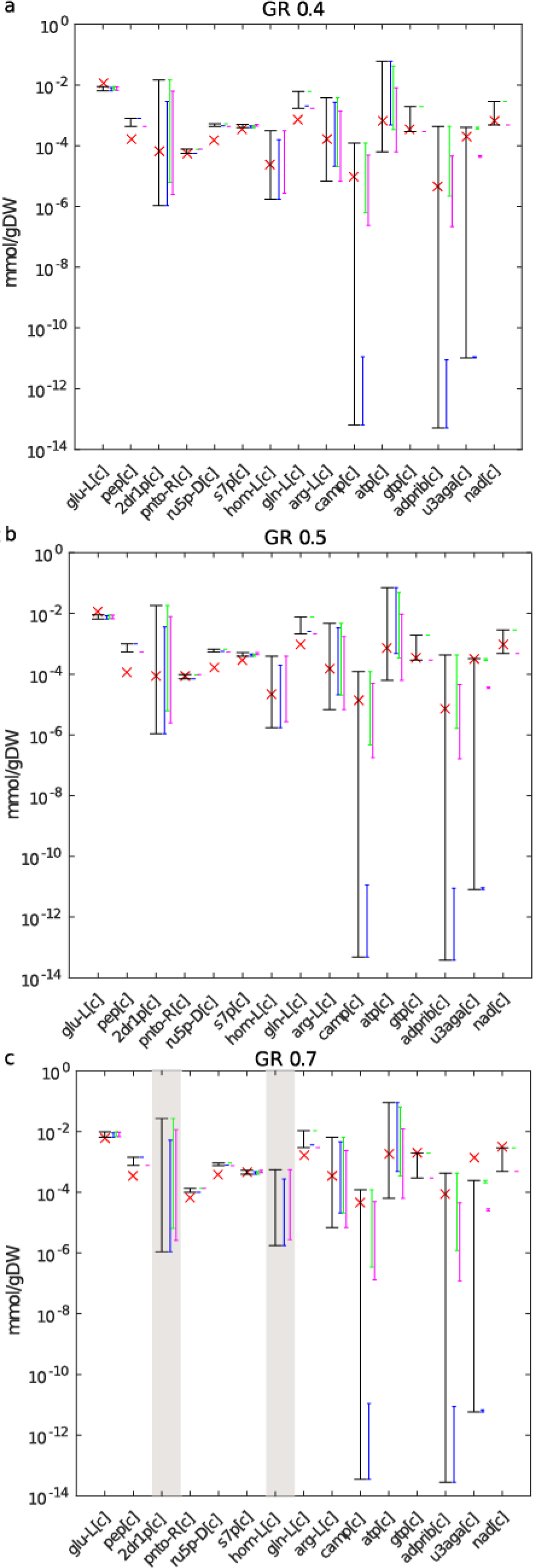
Comparison of predicted ranges with measured metabolite concentrations under the objective of optimizing ATP synthesis and sum of total flux. Comparison of the predicted concentration ranges for 15 intracellular metabolites in *E. coli* with absolute concentrations measured at growth rates (GR) of (**a**) 0.4, (**b**) 0.5 and (**c**) 0.7*h*^−1^. For metabolites with grey background, there is no access to measurements. The colored bars denote the predicted ranges from each of the three different replicates, while the black bar represents the prediction over all replicates. The red cross denotes the measured value at the respective GR. For some metabolites there is no overlap between the colored bars, indicating poor reproducibility over the replicates in the reference scenario. The nomenclature of the metabolites is provided in Supplementary Table S5.

#### Metabolite concentration data set of Gerosa et al. [32]

We use the measurements of steady-state concentrations of 43 metabolites from *E. coli* grown in eight different carbon sources [32]. This data set includes ten of the 199 cytosolic SCC metabolites found in the genome-scale model. We also have access to rates of carbon uptake, some secretion rates, as well as growth from the same experiments (see Supplementary Table S10), which we use as constraints to a genome-scale metabolic network of *E. coli*. Since growth rate is fixed from measurements, as above, we optimize the weighted average of ATP synthesis and total flux, with weighting factors 0.001 for ATP synthesis and 1000 for total flux to reduce the effect of the order difference and make the comparison to optimization of ATP synthesis. Different weighting factors are used in comparison to the analysis of the data set from Ishii et al., above, since different constraints are used that affect the optimal values of the individual objectives. Here, too, at the obtained optimum we can efficiently estimate ranges for the relevant flux ratios (Materials and Methods). To obtain estimates for 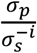, we use the metabolite concentrations from growth on acetate (Supplementary Table S10). We then predict the concentration ranges for the ten SCC metabolites for the seven other carbon sources (Supplementary Fig. S10, S11).

In Supplementary Figures S10 and S11 measured concentration ranges are denoted by red bars and predicted concentration ranges are shown in black. In case of succinate as only carbon source we obtain a model with no feasible solution, so no concentrations could be predicted for that case without further model adaptations. In the remaining growth conditions, depending on the objective and growth condition analyzed, three to five predictions of concentrations resulted in minimum values larger than the respective maximum (missing black bars). This observation is a result of numerical instabilities occurring if flux values *v*_*p*_ and 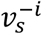 in Eq. (1) differ by several orders of magnitude. The Spearman correlation between the average measured and predicted concentrations (Fig. 5) when optimizing ATP synthesis is 0.63 (p-value 3*10^−4^), while it is only 0.33 (p-value 0.03) when ATP synthesis and total flux are optimized. In addition, the Spearman correlation between the measured and predicted upper and lower bounds when maximization of ATP is used results in higher correlation values (upper bounds 0.61 (p-value 4.3*10^−4^), lower bounds 0.85 (p-value 5.9*10^−9^)) than those when optimization of ATP synthesis and total flux are employed (upper bounds 0.21 (p-value 0.17), lower bounds 0.54 (p-value 1.6*10^−4^)). These findings imply that the usage of different objectives to estimate flux ratios and through them concentrations of metabolites can also be used to discern importance of optimized objectives in a particular experiment.

**Fig. 5.**
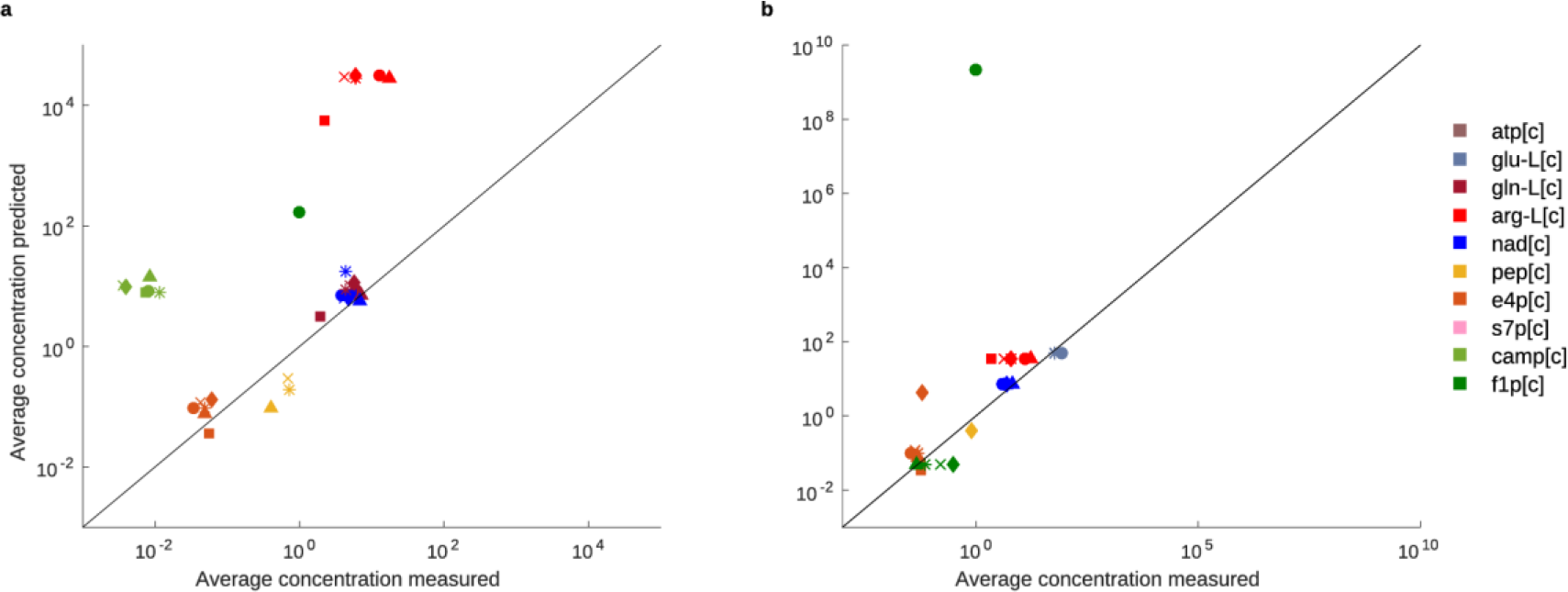
Average measured and predicted concentration of SCC metabolites under different carbon sources. Each data point represents a SCC metabolite (different colors, see legend) under one carbon source (● fructose, ■ galactose, ♦ glucose, ∗ glycerol, × gluconate, ▲ pyruvate). Note that due to numerical instabilities a concentration could not be calculated for all (SCC metabolite, carbon source) combinations, see also Supplementary Fig. S10, S11; concentration prediction using optimization of ATP synthesis and total flux (Spearman correlation 0.33) (b) concentration prediction using optimization of ATP synthesis (Spearman correlation 0.63).

### Changes in metabolite concentrations in knock-out mutants

The fully parameterized kinetic model of *E. coli* can be used to test the applicability of the approach to predict changes in metabolite concentrations in metabolic engineering scenarios. Here, we test the performance of the approach with knock-out mutants based on the following procedure: We make use of the model parameterization to simulate a steady-state concentration and flux distribution from initial physiologically reasonable values for metabolite concentrations. The resulting steady-state concentrations and fluxes yield a wild type reference. We then knock-out each reaction and predict positive steady state flux distribution closest to the wild type reference, following the Minimization of Metabolic Adjustment (MOMA) approach [33]. The resulting flux distribution is used to calculate the concentrations of the 23 SCC metabolites following our approach (Eq. (1)). In the last step, the predicted changes in concentration of the SCC metabolites with respect to the reference are compared to the changes from kinetic simulations of the knock-out with the wild-type reference specifying the initial conditions. We observe similar ranges for the predicted and simulated fold-changes in SCC concentration over all 23 SCC metabolites and knock-outs of 929 reactions for which we were able to simulate a steady-state knock-out flux distribution (Figure 6, fold changes for individual SCC metabolites are shown in Supplementary Figure 12). We grouped the fold-changes into 12 bins, given in the x-axis of Figure 6. For ten SCC metabolites, the predicted fold change of at least 29% of the knock-outs is in the same bin as the simulated fold change. The highest overlaps are observed for AMP (39%), phosphoenolpyruvate (38%) and isocitrate (37%). In contrast, the fold changes in concentration for metabolites like succinyl-CoA, acetyl-CoA, oxaloacetate, malate and pyruvate are in the same class as simulated for at most 1% of the knock-outs. The lack of correspondence between simulated and predicted concentrations for some SCC metabolites (Supplementary Fig. S12) indicates that principles others than those used in MOMA shape the metabolic adjustment of knock-out mutants.

**Fig. 6.**
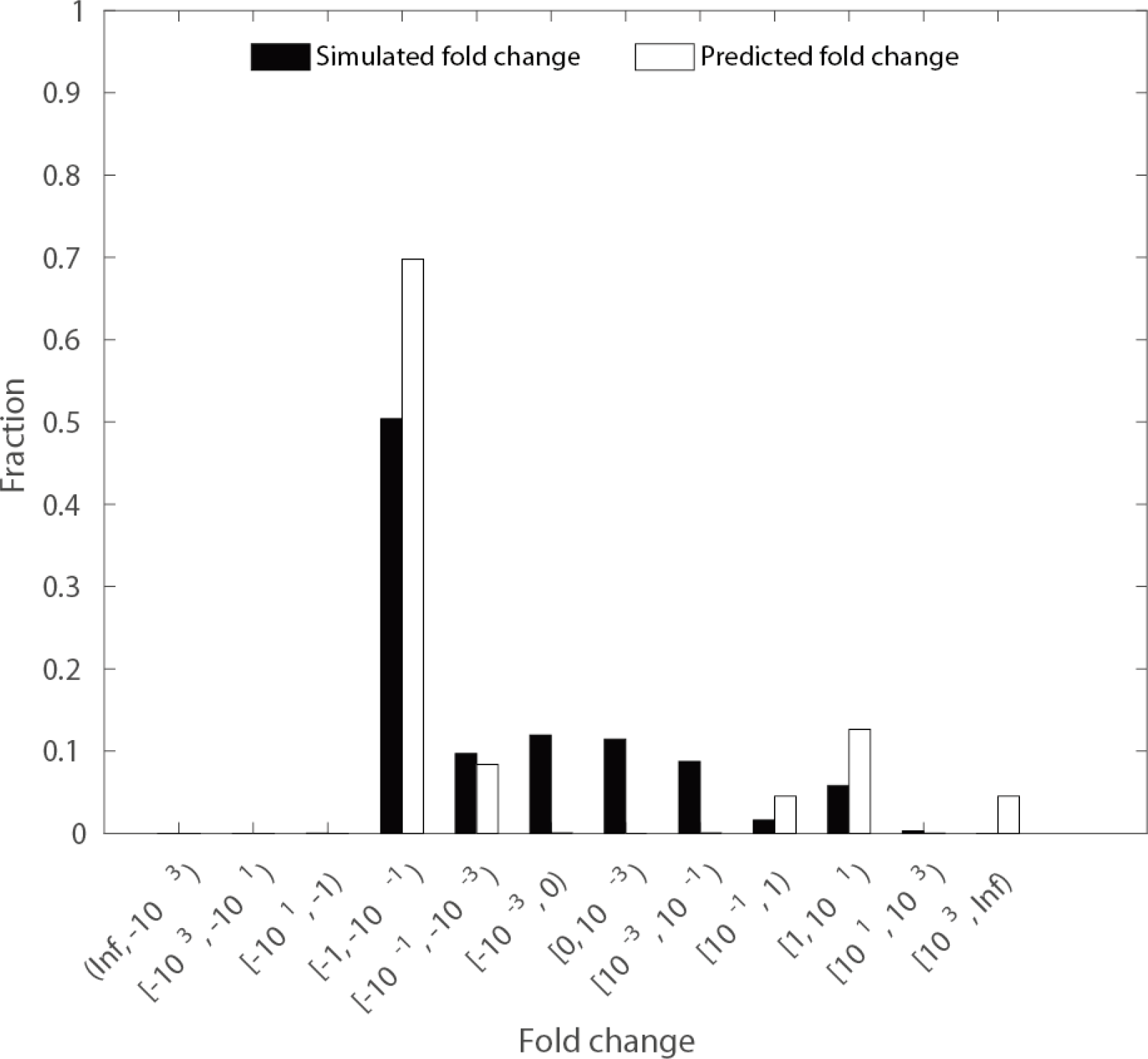
Fold change in concentration of SCC metabolites upon reaction knock-out. The distribution of predicted and simulated fold change in concentration of 23 SCC metabolites over 929 single knock-out mutants for which a steady-state flux distribution could be simulated.

### Metabolites with SCC across species

We next apply Eq. (1) to 14 large-scale metabolic networks which differ in complexity due to the number of considered metabolites and reactions as well as their organization in subcellular compartments (Supplementary Table S6). The investigated metabolic networks are mass-and charge-balanced and support positive steady-state reaction rates (see Methods). Since reliable kinetic information and measurements of absolute concentration measurements are currently missing across diverse species, we report only the number of the metabolites with SCC across the analyzed large-scale networks.

We find that the percentage of metabolites with SCC ranges from 7.74 % and 8.02% in the models of *N. pharaonis* and *C. reinhardtii* to 33.66% and 36.53% in the models of *A. thaliana* and *Y. pestis* (Fig. 7a). Interestingly, the number of metabolites with SCC scales linearly with the total number of metabolites (Fig. 7b, *R*^2^ = 0.82) and the number of reactions in the examined networks (Fig. 7c, *R*^2^ = 0.76). This finding indicates that the proposed approach is not limited to networks of a particular size.

**Fig. 7.**
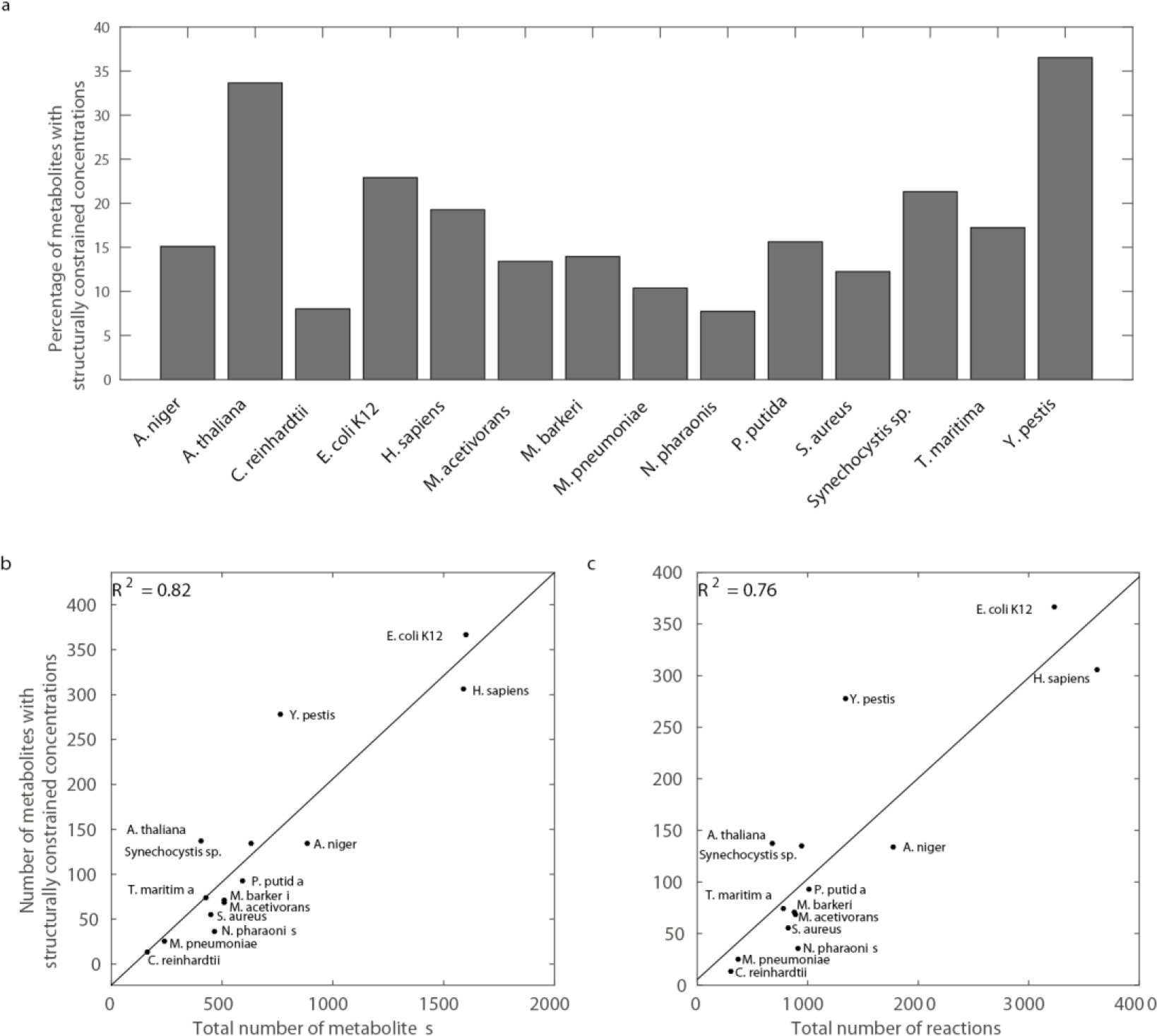
Metabolites with structurally constrained concentration across species. (a) The fraction of metabolites with structurally constrained concentrations in 14 large-scale metabolic networks from all kingdoms of life. The number of these metabolites scales linearly with (b) the total number of metabolites (*R*^2^=0.82) and (c) the total number of reactions (*R*^2^=0.76).

Different reasons can be used to explain the observation that larger networks contain more metabolites with SCC. For instance, larger networks may include more linear pathways, whereby the number of reactions which are fully coupled due to structure is expected to increase. Yet, in denser networks, which include more reactions on the same set of metabolites, it is more likely to identify reactions which share substrates of same stoichiometry, which then leads to full coupling due to mass action kinetics, as considered in our approach. To investigate the reasons for the scaling of the number of metabolites with SCC, we determine the number of: (*i*) metabolites which are synthesized and used by one reaction, respectively (in support of the linear pathway explanation), (*ii*) fully coupled reactions only due to structure, (*iii*) coupled reactions due to mass action (in support of the network density explanation), (*iv*) the combination of (*ii*) and (*iii*), to assess the couplings due to both structure and kinetics (Supplementary Table S7). We calculate the Pearson correlation coefficient between each of these properties and the number of reactions over the analyzed networks, as a measure of network size (Supplementary Table S7). Larger networks indeed contain a bigger number of metabolites synthetized and used by a single reaction, respectively, and more reactions which are fully coupled due to both structure and kinetics. Therefore, both the linear pathway and the network density explanations contribute to the observed scaling in the analyzed networks.

Due to the derivation of Eq. (1), it may be expected that the approach is not applicable to metabolites which participate in a large number of reactions, since they may be less likely to be fully coupled. Nevertheless, our findings show that between 28.89% and 62.95% of the SCC metabolites in the analyzed networks are involved in more than two reactions (see Supplementary Table S6). One reason is that a SCC metabolite may also be determined by applying Eq. (1) to the ODE of another metabolite (see Eq. (2) and Fig. 1c).

Since changes in relevant fluxes directly affect the concentration of a SCC metabolite, they can be used to tightly control the concentration range. For essential metabolic processes to be carried out efficiently, metabolites that serve as coenzymes and energy currency of biological systems, namely, the oxidized and reduced version of NAD and NADP as well as the adenosine phosphates (i.e. AMP, ADP, ATP), are maintained within certain concentration ranges that can be readily controlled, as is the case for SCC metabolites. Despite the many biochemical reactions in which these ubiquitous metabolites participate (Supplementary Table S8), all of which must satisfy our conditions in order to invoke Eq. (1), we find that the (sub)cellular concentrations of ATP and NAD are indeed structurally constrained in twelve and ten of the analyzed networks, respectively. This implies that the network structure, alongside the relevant rate constants and relevant flux ratio, imposes boundaries on and facilitates simple control over their concentrations. In addition, we find that NADP shows SCC in four of the investigated networks, including *A. thaliana* and *C. reinhardtii* (Table 1 and Supplementary Table S8). In these photosynthetic organisms, NADPH is produced by ferredoxin-NADP+ reductase in the last step of the electron transport chain which constitutes the light reactions of photosynthesis [34]. The produced NADPH provides reducing power for the biosynthetic reactions in the Calvin cycle to fix carbon dioxide as well as in the reduction of nitrate into ammonia for plant assimilation in the nitrogen cycle. Therefore, precise and simple control of NADPH will provide uninterrupted functionality of these key metabolic pathways and maintenance of carbon and nitrogen balance [35]. In addition, for ten models, we find that H+ is SCC, ensuring maintenance of the specific functions of individual organelles [36]. Altogether, our findings indicate that the concentration ranges for coenzymes and other components essential for fueling metabolism can be established by controlling few ratios of fluxes, despite their involvement in hundreds of reactions. Moreover, they imply that the network architecture may be organized such that the concentrations of these metabolites are intrinsically constrained and easy to control.

**Table 1.**
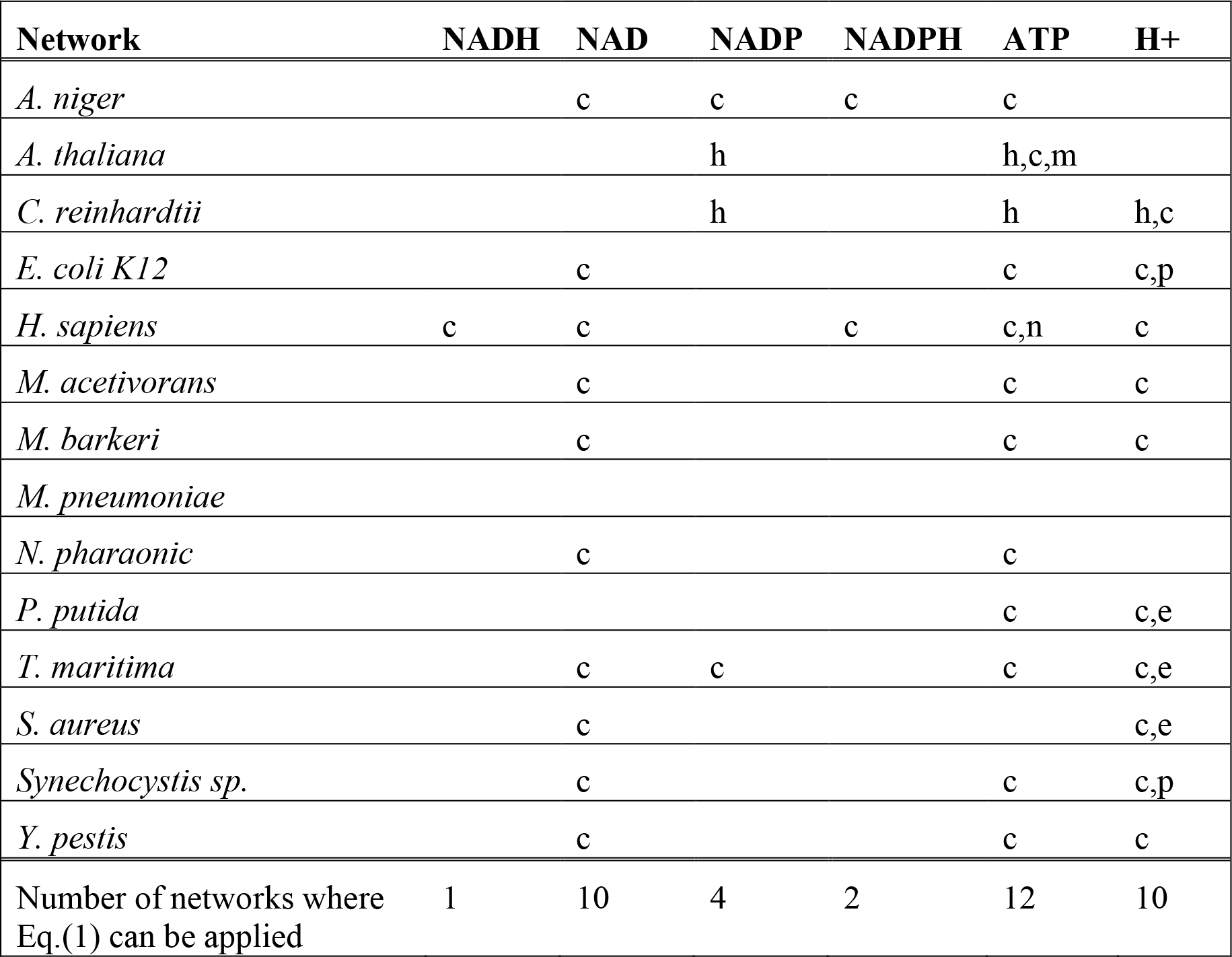
Structurally constrained concentrations for metabolites serving as energy currency. (h=chloroplast, c = cytosol, m = mitochondria, n = nucleus, p = periplasm, e = extracellular space). The table summarizes the networks in which Eq. (1) holds for NADH, NAD, NADP, NADPH, ATP, and H+. The table includes the respective compartments in which Eq. (1) can be applied for the investigated metabolites.

## Discussion

Genome-scale metabolic networks have propelled the understanding of the metabolic capabilities for a wide variety of organisms across all kingdoms of life. The existing large-scale modelling approaches examine the space of feasible fluxes, but cannot be used to infer the metabolite concentrations driving these fluxes without extensively relying on largely unknown kinetic parameters. Hence, the direct usage of large-scale metabolic networks to make predictions about concentrations that are directly testable from high-throughput metabolomics data is not possible with the existing modelling approaches.

Here we derive a condition that pinpoints that the structure of a metabolic network, ratios of relevant rate constants, and ratios of relevant reaction fluxes constitute the determinant of concentration ranges for selected metabolites. This link is based on the well-known concept of full coupling of reactions [21, 23] which we expand under the assumption of mass action kinetics to include reactions that share substrates of same stoichiometry. These concepts allow us to efficiently determine the admissible concentration ranges in large-scale metabolic networks endowed with mass action kinetics across all kingdoms of life. The derivation of Eq. can be generalized by considering reactions which differ in order larger than one with respect to a single metabolite. For a given flux distribution this approach results in a polynomial equation in a single variable which can be efficiently solved with the Newton’s method.

Our approach is also applicable to networks with kinetic laws derived from mass action which involve enzyme forms (e.g., Michaelis-Menten). This can be achieved by augmenting the network to include reactions which model substrate-enzyme complex formation as well as the synthesis and degradation of enzymes. However, these extensions come at a cost of substantially larger data sets which are not yet readily available. In addition, our analyses demonstrate that the casting of a kinetic rate law in terms of mass action mechanisms may affect the findings regarding the SCC metabolites. For instance, we find that there are many more SCC metabolites in comparison to other SCC components (i.e., enzymes and enzyme-substrate complexes) in each of the analyzed models (Supplementary Table S9). With exception of the network of *C. reinhardtii*, the usage of enzymatic forms explicitly in mass action mechanisms leads to a decrease in the number of metabolites with SCC (Supplementary Table S9), due to the decrease in the number of reaction pairs which differ in their order by one. Applications of the approach to other forms of kinetics will be subject in future investigations and extensions.

Our approach provides a links between metabolite concentrations, relevant rate constants, and relevant flux ratios; therefore, information on two of these can be used to predict the third. Our analyses demonstrate that there is a good quantitative agreement between predicted and simulated concentration ranges based on full knowledge of rate constants from a kinetic model of *E. coli*. Rate constants of elementary reactions are expected to become increasingly available for model organisms, largely due to the development of computational methods coupled with high-throughput data [8, 9]. In addition, by examining the scenario where flux ratios are estimated from the constraint-based modeling framework, we observe that the approach can be used to select which objective function (or a combination thereof) is optimized by a biological system for which metabolite concentration measurements are available.

Most importantly, we show that even in the absence of data on relevant rate constants and relevant flux ratios, we can apply the approach to successfully predict concentration ranges in *E. coli* under different growth conditions, provided measurements of concentrations for SCC metabolites in one reference condition. Therefore, the proposed approach represents an important step in complementing genome-scale metabolic networks with metabolite concentrations, widening the applicability of large-scale models to a range of biotechnological and medical applications.

## Materials and Methods

### Components with structurally constrained concentrations

A metabolic network can be represented by the stoichiometric matrix, *N* = *N*^+^ − *N*^−^, where *N*^+^ includes the stoichiometry of the products and *N*^−^ comprises the stoichiometry of the substrates of each reaction. In the following, we derive the conditions for structurally constrained robustness of component *X*_*i*_ based on the ordinary differential equation (ODE) for the component *X*_*j*_ (not necessarily different from *X*_*i*_) under the assumption that the reaction rates, *v*(*t*), satisfy mass action kinetics, whereby 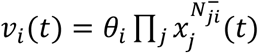. Let the ODE be specified by 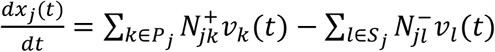, where *P*_*j*_ is the set of reactions with *X*_*j*_ as one of their products and *S*_*j*_ is the set of reactions which have metabolite *X*_*j*_ as one of their substrates.

We consider the following two cases: (*i*) the concentration of *X*_*i*_ appears in every *v*_*k*_(*t*) for which 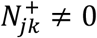 and for every *v*_*k*_(*t*) there exist a set 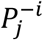 of reactions 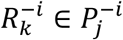 such that 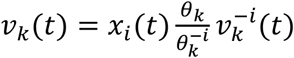 and (*ii*) the concentration of *X*_*i*_ appears in every *v*_*l*_(*t*) for which 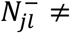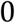 and for every *v*_*l*_(*t*) there exist a set of reactions 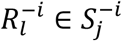 such that 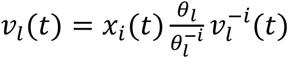.

#### Case I

The rates of a reaction *R*_*k*_ and a reaction from the set 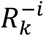 are given by 

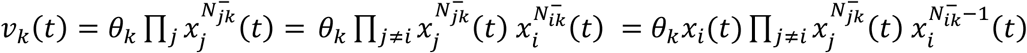

and 
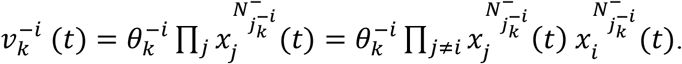

From rewriting the equation of 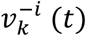 above we have that 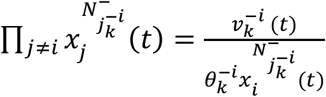. Since 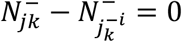 for every *j* ≠ *i* and 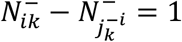 we can rewrite the equation of *v*_*k*_(*t*) such that 

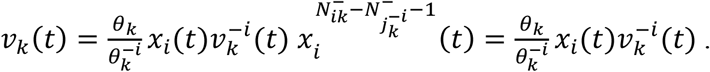

The ODE for component *X*_*j*_ revealing structurally constrained concentration of component *X*_*i*_ is then given by: 

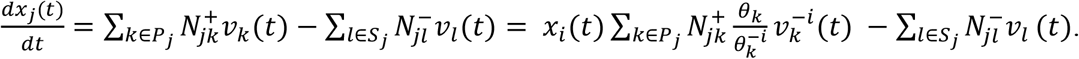

Let *p* and *s* bet two reaction indices such that 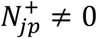 and 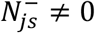. In any positive state *v*(*t*), we have that 

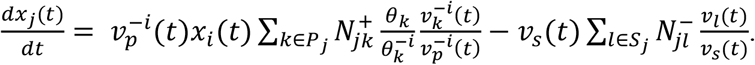

In a steady state then 

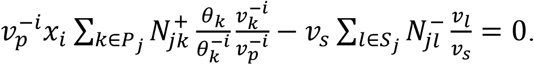

If for every 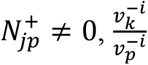 is constant because either reactions 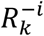 and 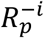 are fully coupled or share the same substrates, then 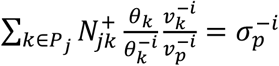 is a constant that only depends on a subset of rate constants and the network structure. Moreover, if for every 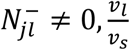 is constant because either reactions *R*_*l*_ and *R*_*s*_ are fully coupled or share the same substrates, then 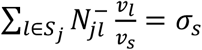 is a constant, too, which in the simplest case when all reactions in *S*_*j*_ are fully coupled irrespective of the kinetic rate law, only depends on the network structure. Therefore, 

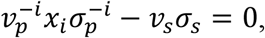

and 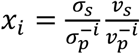

For each reaction *R*_*k*_ in *S*_*j*_, there exists a non-empty subset 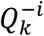 of reactions lacking one substrate molecule of *X*_*i*_ in comparison to *R*_*k*_; the union of all 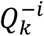 yields the set of reactions 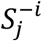. Let *Q* be a subset of 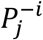 that contains one and only one reaction from each of 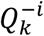. Since the reaction indices *p* and *s* are arbitrarily chosen, the concentration range of metabolite *X*_*i*_ for a given subset *Q* over a given set of flux distributions, *F*, is given as 

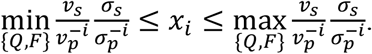

#### Case II

The rates of a reaction *R*_*l*_ and a reaction from the set 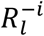 are given by 

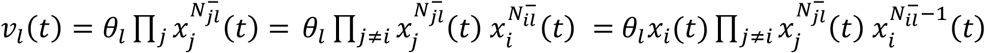

and 
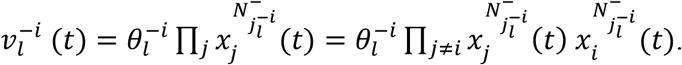

From rewriting the equation of 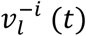 above we have that 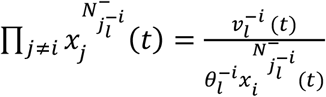. Since 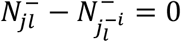 for every *j* ≠ *i* and 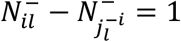 we can rewrite the equation of *v*_l_ (*t*) such that 

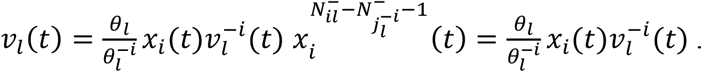

The ODE for component *X*_*j*_ revealing structurally constrained concentration of component *X*_*i*_ is then given by: 

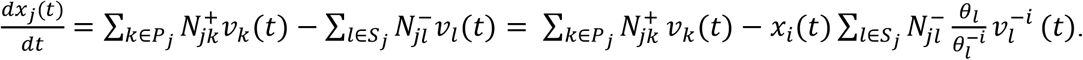

Let *p* and *s* bet two reaction indices such that 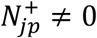 and 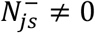. In any positive state *v*(*t*), we have that 

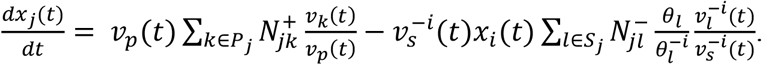

In a steady state then 

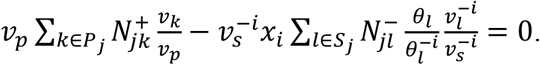

If for every 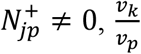 is constant because either reactions *R*_*k*_ and *R*_*p*_ are fully coupled or share the same substrates, then 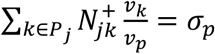 is a constant that, in the simplest case when all reactions in *P*_*j*_ are fully coupled irrespective of the kinetic rate law, depends only on the network structure. Moreover, if for every 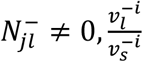 is constant because either reactions 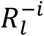 and 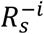 are fully coupled or share the same substrates, then 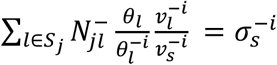. The constant 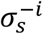 then only depends on a subset of rate constants and the network structure.

Therefore, 

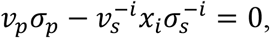

and 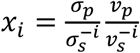

For each reaction *R*_*l*_ in *P*_*j*_, there exists a non-empty subset 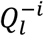 of reactions lacking one substrate molecule of *X*_*i*_ in comparison to *R*_*l*_; the union of all 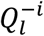 yields the set of reactions 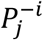. Let *Q* be a subset of 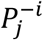 that contains one and only one reaction from each of 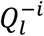. Since the reaction indices *p* and *s* are arbitrarily chosen, the concentration range of metabolite *X*_*i*_ for a given subset *Q* over a given set of flux distributions, *F*, is given as 

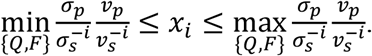

As a result, the ranges for steady-state concentration *x*_*i*_ can be expressed as a function of a set of given flux distributions, ratios of specific fluxes and constants that depend only on the structure of the network and values for a subset of rate constants. Since fluxes are the integrated outcome of transcription, translation, and post-translational modifications and their interplay with the environment and nutrient availability, our derivation provides a direct relation between concentration ranges, flux ratios, and rate constants.

### Flux coupling

Let *C*(*N*) = {*v* ∈ ℝ^*n*^|*Nv* = 0, *v* ≥ 0} be the steady-state flux cone for a given stoichiometric matrix *N* with *n* reactions, under the assumption that every reaction is irreversible. Here, we restrict our analysis to the subspace *F* ⊂ *C*(*N*) by bounding the fluxes: *F* = {*v* ∈ ℝ^*n*^|*N v* = 0,0 ≤ *lb* ≤ *v* ≤ *ub*}, where *lb* and *ub* are lower and upper flux bounds. We will refer to *v* ∈ *F* as the feasible flux distributions. A reaction *R*_*i*_ is called blocked if for every *v* ∈ *F*, *v*_*i*_ = 0. A pair of reactions *R*_*i*_ and *R*_*j*_ is called fully coupled, if there exists *λ* > 0, such that for every *v* ∈ *F*, *v*_*i*_ = *λv*_*j*_

The minimum and maximum value for the ratio 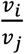 over the flux distributions in *F* can be determined by the linear-fractional programming: 

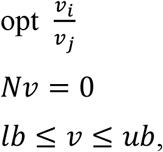

which can be rewritten following the Charnes-Cooper transformation [37] to the following linear program: 

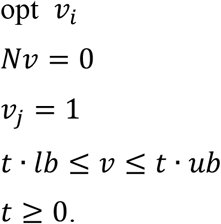

If the minimum and maximum values for the linear program are the same, then the reactions *R*_*i*_ and *R*_*j*_ are fully coupled. Such reactions can be efficiently computed for large-scale networks[4, 21].

In addition, under the mass action kinetics, two reactions are fully coupled in any state of the system if they share the same substrates with the same stoichiometry. This leads to additional full couplings due to the transitivity of the relations, as demonstrated in the main text.

### Metabolites with structurally constrained concentrations in mass action networks

In the following, we present an algorithm determining SCC metabolites under the assumption of mass action kinetics:

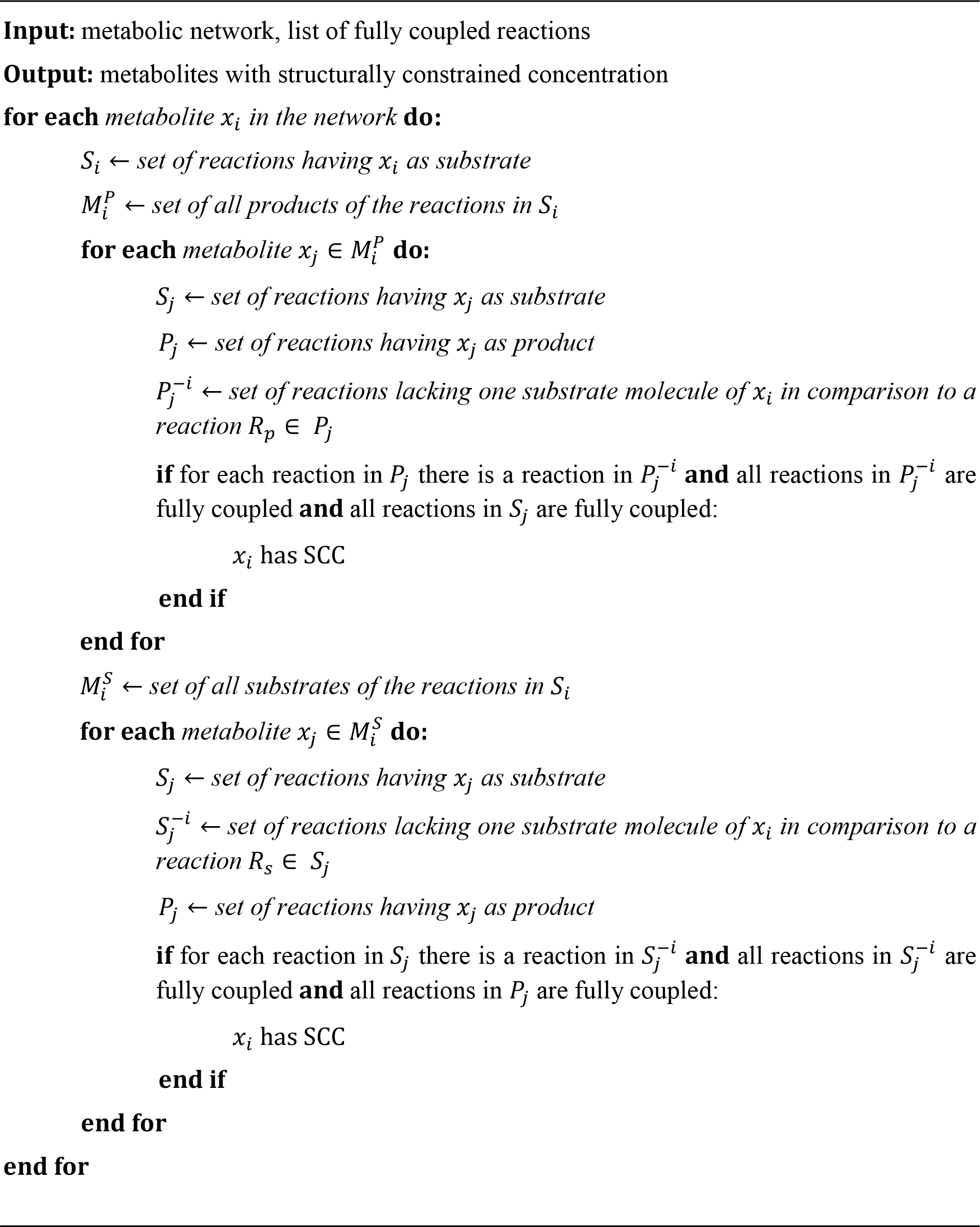

### Correlation analysis

Using a large-scale kinetic model of *E. coli* we simulate 100 steady-state flux distribution and steady-state concentrations from different initial concentrations. Initial concentrations were obtained by perturbation of the original initial concentration of a metabolite by 1, 5, 10 or 20%. We run the model until a steady-state was reached. Using the simulated steady-state flux distributions we can predict concentration ranges for 23 metabolites using Eq. (2) (Supplementary Table S1). The Pearson correlation was then calculated for *(i)* simulated and predicted upper bounds, *(ii)* simulated and predicted lower bounds, and *(iii)* the absolute range over simulated and predicted concentrations. In addition, we also determined the correlation between shadow price for the respective metabolites and the simulated range, as well as, to the coefficient of variation obtained over simulated concentrations (Supplementary Table S2). Moreover, we calculated the Euclidean distance between upper and lower bound from prediction and simulation, respectively. Due to the high difference in the order of magnitude over the analyzed metabolites we also calculated Euclidean distance after normalizing the data. We considered the Euclidean distance of log-transformed concentration vectors, and the Euclidean distance between the concentration vectors normalized by the respective maximum value.

### Effect of missing information on rate constants

To assess the effect of missing information about rate constants on the accuracy of the predicted concentration range, we simulated missing knowledge about parameters by removing 10, 30, 50, 70 or 90% of the relevant rate constants uniformly at random. We consider only removing information about relevant rate constants to avoid bias due to removal of information in parts of the network that have no effect on the predictions of the concentration ranges. We compare the Pearson and Spearman correlation coefficient between predicted and simulated concentration ranges as well as the two versions of Euclidean distance for each percentage obtained over 100 random removals of rate constants.

### Effect of missing information on flux ratios

To assess the effect of missing information about flux ratios on the accuracy of the predicted concentration range, we obtained relevant flux ratios from constraint-based modeling. Therefore, we solve the following linear program optimizing a weighted average of ATP production and total flux: 

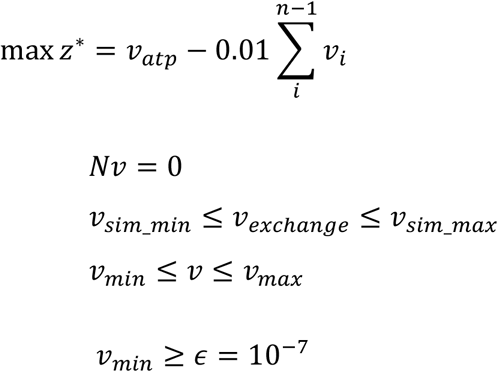

In addition, the flux through exchange reactions is constrained by the respective minimum, *v*_*sim_min*_, and maximum value, *v*_*sim_max*_, obtained over 100 simulations (Supplementary Table S1) to obtain a physiologically reasonable flux distribution. The weighting factor of 0.01 was chosen to reduce the effect of three orders of magnitude difference in the respective optimum observed when ATP production and total flux are optimized individually.

Next, we determine the range for the relevant flux ratio 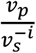 at the optimum *z*\* using a transformed linear-fractional program: 

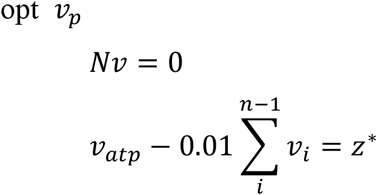

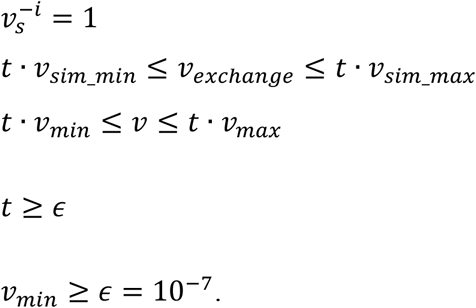

We then used the obtained ranges for 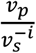 together with Eq. (2) to calculate concentration ranges for SCC metabolite *X*_*i*_.

### Extension of the approach based on available concentration measurements

Using the most recent genome-scale metabolic network of *E. coli* [27] together with measurements of steady-state concentrations from *E. coli* under different growth scenarios [28] we predict concentration ranges for 15 SCC metabolites using the following procedure. We first use the concentration measurements from three replicates at a growth rate of 0.2*h*^−1^ (reference state) together with flux ratios obtained from constraint-based modelling to estimate the ratio 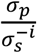 given that 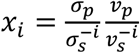

For each replicate we solve the following linear programs in order to obtain ranges for the relevant flux ratios 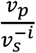. 

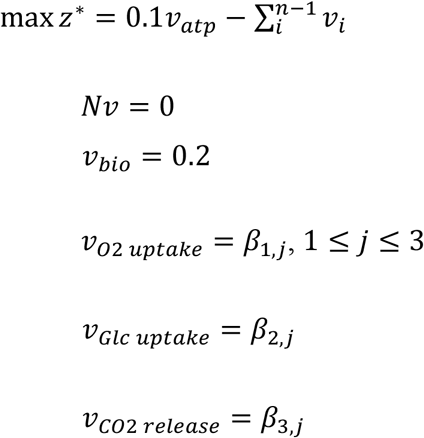

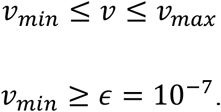

The linear program above constrains rates of glucose and oxygen uptakes, carbon dioxide release as well as growth by values *β*_*i,j*_ (which differ between replicates *j*, 1 ≤ *j* ≤ 3) available from measurements [28]. We optimize the weighted average of ATP synthesis and total flux. The weighting factor of 0.1 and 0.001 for ATP synthesis, for the data set of Ishii et al. [28] and Gerosa et al. [32], respectively, is chosen to reduce the effect of the order difference in the respective optimum observed when ATP production and total flux are optimized individually. In addition, we use weighting factors of 1 and 1000 for optimization of total flux in the case of Ishii et al. [28] and Gerosa et al. [32], respectively. To obtain ranges for the relevant flux ratios 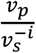, which are employed to calculate ranges for ratios 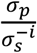, we solve the following linear program at the optimum *z*\*: 

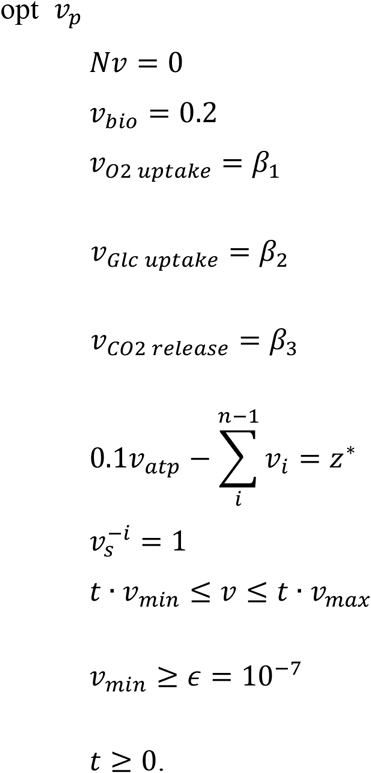

From Eq. (2) we predict concentration values for *E. coli* cells with growth rates of 0.4, 0.5, and 0.7*h*^−1^ using the previously obtained estimates for ranges of 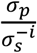 together with ranges of 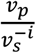. The latter can be obtained following the same procedure as described above using rates of glucose and oxygen uptakes, carbon dioxide release as well as growth for *E. coli* cells grown at rates of 0.4, 0.5, and 0.7*h*^−1^.

### Fold changes in SCC metabolite concentrations in knock-out mutants

We use a large-scale kinetic model of *E. coli* [8] to simulate a steady-state concentration and flux distribution from initial physiologically reasonable values for metabolite concentrations provided in the original publication. The simulated steady-state concentrations and fluxes yield a wild type reference. Next, we simulate single reaction knock-outs and predict positive steady state flux distribution closest to the wild type reference, following the Minimization of Metabolic Adjustment (MOMA) approach [33] for each mutant. The resulting flux distribution is used to calculate the concentrations of the 23 SCC metabolites following Eq. (1). In addition, we simulate steady-state flux distributions and concentrations for knock-out mutants from the kinetic model using the wild type reference as initial concentrations. For 929 out of 1474 reaction knock-outs we could simulate steady-state values. Based on these knock-out mutants we then compare fold changes in concentration of the SCC metabolites with respect to the reference obtained from kinetic model simulations and predictions using MOMA.

## Acknowledgements

J.M.O.E.M., A.K., G.B., and Z.N. are supported by the Max Planck Society. The authors and the Max Planck Society have initiated a process for patenting the computational approach and findings presented in this manuscript.

## Author contributions

Conceptualization, Z.N.; Methodology, J.M.O.E.-M., Z.N.; Software, A.K.; Formal Analysis, A.K, Z.N.; Investigation, A.K., G.B.; Writing - Original Draft, G.B., Z.N.; Writing - Review and Editing, A.K., J.M.O.E.-M., G.B., and Z.N.

## Competing interests

The authors declare no competing interests.

## Data availability

The code and data to reproduce the results are available on GitHub https://github.com/ankueken/SCC.

## Supplementary Figure legends

**Fig. S1. Agreement between simulated and predicted bounds from a kinetic metabolic model of *E. coli***. The simulated and predicted (**a**) lower and (**b**) upper concentration bounds for 23 SCC metabolites in the large-scale kinetic model of *E. coli*. The very small discrepancies are due to numerical instabilities.

**Fig. S2. Distribution of rate constants used in calculation of concentration ranges for SCC metabolites in a genome-scale metabolic model of *E. coli***. Distribution of (**a**) the relevant rate constants and (**b**) their ratios for reactions coupled due to mass action kinetics; log-log distribution of (**c**) the relevant rate constants and (**d**) their ratios for reactions coupled due to mass action kinetics.

**Fig. S3. Effect of missing information about relevant rate constants on the accuracy of concentration range predictions for a large-scale kinetic model of *E. coli*.** We consider 10–90% of the relevant rate constants to be unknown by random removal. We consider three scenarios for the substitution of missing ratios of rate constants: (*i*) equality (i.e., kinetic rate constants are assumed to be the same), (*ii*) the mean, or (*iii*) the median of the ratios of relevant rate constants that are still present in the model. Shown are the boxplots (red lines inside each box denote the corresponding medians) of the resulting Spearman correlation coefficients between the predicted and simulated (**a**) lower bound vectors and (**b**) upper bound vectors of concentrations over the SCC metabolites in the kinetic model of *E. coli*.

**Fig. S4. Effect of missing information about relevant rate constants on the accuracy of concentration range predictions for a large-scale kinetic model of *E. coli***. We consider 10–90% of the relevant rate constants to be unknown by random removal. We consider three scenarios for the substitution of missing ratios of rate constants: (*i*) equality (i.e., kinetic rate constants are assumed to be the same), (*ii*) the mean, or (*iii*) the median of the ratios of relevant rate constants that are still present in the model. Shown are the boxplots (red lines inside each box denote the corresponding medians) of the average Euclidean distance between the predicted and simulated (**a**) lower bound vectors and (**b**) upper bound vectors of concentrations over the SCC metabolites in the kinetic model of *E. coli*.

**Fig. S5. Effect of missing information about relevant rate constants on the accuracy of concentration range predictions for a large-scale kinetic model of *E. coli***. We consider 10–90% of the relevant rate constants to be unknown by random removal. We consider three scenarios for the substitution of missing ratios of rate constants: (*i*) equality (i.e., kinetic rate constants are assumed to be the same), (*ii*) the mean, or (*iii*) the median of the ratios of relevant rate constants that are still present in the model. Shown are the boxplots (red lines inside each box denote the corresponding medians) of the Euclidean distance between the log-transformed predicted and log-transformed simulated (**a**) lower bound vectors and (**b**) upper bound vectors of concentrations over the SCC metabolites in the kinetic model of *E. coli*.

**Fig. S6. Effect of missing information about relevant rate constants on the accuracy of concentration range predictions for a large-scale kinetic model of *E. coli***. We consider 10–90% of the relevant rate constants to be unknown by random removal. We consider three scenarios for the substitution of missing ratios of rate constants: (*i*) equality (i.e., kinetic rate constants are assumed to be the same), (*ii*) the mean, or (*iii*) the median of the ratios of relevant rate constants that are still present in the model. Shown are the boxplots (red lines inside each box denote the corresponding medians) of the Euclidean distance between the predicted and simulated (**a**) lower bound vectors of concentrations normalized by the respective maximum value and (**b**) upper bound vectors of concentrations normalized by the respective maximum value over the SCC metabolites in the kinetic model of *E. coli*.

**Fig. S7. Predicted concentration ranges for 15 intracellular metabolites in *E. coli* at growth rates (GR) of 0.4, 0.5 and 0.7 *h*^−1^ under the objective of optimizing ATP synthesis and sum of total flux.** The bars denote the predicted ranges from each of the three different scenarios (**a**) over all three replicates and (**b**) over replicates with not more than one magnitude difference in estimated range for the ratio of 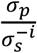The marked points denote the measured concentrations in the employed data set.

**Fig. S8. Distribution of average Euclidean distance between simulated and predicted concentration.** From each of the 100 simulated steady-state flux distributions we predict concentrations for the SCC metabolites and calculate the average Euclidean distance between the simulated and predicted concentrations.

**Fig. S9. Comparison of predicted ranges with measured metabolite concentrations under the objective of optimizing ATP synthesis for the data set of Ishii et al.** Comparison of the predicted concentration ranges for 15 intracellular metabolites in *E. coli* with absolute concentrations measured at growth rates (GR) of (**a**) 0.4, (**b**) 0.5 and (**c**) 0.7*h*^−1^. The colored bars denote the predicted ranges from each of the three different replicates, while the black bar represents the prediction over all replicates. For some metabolites no value could be predicted due to numerical instabilities. The red cross denotes the measured value at the respective GR. For metabolites with missing red cross, there is no access to measurements. The nomenclature of the metabolites is provided in Supplementary Table S5.

**Fig. S10. Comparison of predicted ranges with measured metabolite concentrations under the objective of optimizing ATP synthesis and total flux for the data set of Gerosa et al.** Comparison of the predicted concentration ranges for 10 intracellular metabolites in *E. coli* with absolute concentrations measured at seven different carbon sources. The red bars denote the measured ranges over three different replicates, while the black bar represents the predicted concentration. For some metabolites no value could be predicted due to numerical instabilities. For the model simulating growth on succinate no steady-state solution could be obtained without further model adaptation, therefore, no SCC concentration could be predicted.

**Fig. S11. Comparison of predicted ranges with measured metabolite concentrations under the objective of optimizing ATP synthesis for the data set of Gerosa et al.** Comparison of the predicted concentration ranges for 10 intracellular metabolites in *E. coli* with absolute concentrations measured at seven different carbon sources. The red bars denote the measured ranges over three different replicates, while the black bar represents the predicted concentration. For some metabolites no value could be predicted due to numerical instabilities. For the model simulating growth on succinate no steady-state solution could be obtained without further model adaptation, therefore, no SCC concentration could be predicted.

**Fig. S12. Fold change in concentration of SCC metabolites upon reaction knock-out.** Distributions of predicted and simulated fold change in concentration for the 23 SCC metabolites over 929 single knock-out mutants, for which a steady-state flux distribution could be simulated.

### Supplementary Table captions

**Table S1. (A)** Initial conditions sampled for simulations of the large-scale kinetic model of *E. coli*. The initial concentration is given in units mmol/gDW. (**B**) Steady-state concentrations obtained from simulations of the large-scale kinetic model of *E. coli* starting from the respective initial conditions presented in Table S1A. The first two columns show the respective minimum and maximum steady-state concentration over all 100 simulations. The concentration is given in units mmol/gDW. (**C**) Steady-state flux distributions obtained from simulations of the large-scale kinetic model of *E. coli* starting from the respective initial conditions presented in Table S1A. The flux is given in units mmol/gDW/hr. (**D**) Simulated and predicted concentration ranges for 23 SCC metabolites in a kinetic metabolic model of *E. coli*.

**Table S2.** (**A**) Correlation between predicted concentration range and shadow price for 23 structurally constrained metabolites to the corresponding metabolic concentrations obtained from 100 simulations of a kinetic model of *E. coli* core metabolism. (**B**) Euclidean distance between simulated and predicted concentration bounds for 23 SCC metabolites in large-scale kinetic model of *E. coli*. In addition the table provides simulated and predicted concentration bounds in mmol/gDW.

**Table S3.** List of rate constants for reactions in the genome-scale model iJO1366 of *E. coli*. In addition to the used rate constants and the related organism in BRENDA, the table reports the reaction abbreviation used in the model and the enzyme EC number related to each reaction. In case more than one rate constant is known per reaction we consider the average value.

**Table S4.** (**A**) Measured concentrations of SCC metabolites in *E. coli* under different growth scenarios. The three replicates at growth rate 0.2h^−1^ are used as reference state. Measured volumetric concentrations^1^ were converted to mmol/gDW by using a ratio of aqueous *E. coli* cell volume to dry weight of 0.0023L/g^2^. (**B**) Specific flux rates for *E. coli* grown under different scenarios.

**Table S5.** (**A**) Predicted concentration ranges for the 15 SCC metabolites in a genome-scale metabolic model of *E. coli* with available data on concentration. (**B**) In addition correlation values between predicted and simulated bounds are provided.

**Table S6.** Number of metabolites with structurally constrained concentrations for each of the metabolic networks analyzed. The numbers of reactions and metabolites correspond to the number after reaction splitting into irreversible reactions and removal of blocked reactions. The latter is needed to satisfy the prerequisite for a positive steady state.

**Table S7.** Fraction of fully coupled reactions and reactions coupled due to mass action kinetics in 14 analyzed genome-scale metabolic networks.

**Table S8.** Structurally constrained metabolites across the 14 analyzed metabolic networks. In addition, the in-and out-degree for these metabolites are provided. Metabolites marked in red correspond to energy metabolism (see Table 1 in the main text) and metabolites marked in green exhibit absolute concentration robustness. Metabolite names and their abbreviations are used as provided in the original models.

**Table S9.** Number of metabolites with structurally constrained concentrations metabolic networks analyzed including enzyme information. The numbers of reactions and metabolites correspond to the number after rewriting in Michaelis-Menten format, reaction splitting into irreversible reactions and removal of blocked reactions. Model components correspond to metabolites, enzymes and enzyme-substrate-complexes.

**Table S10.** (**A**) Measured concentrations of SCC metabolites in E. coli under growth on different carbon sources. Replicates for growth on acetate are used as reference state. (**B**) Specific flux rates for E. coli under growth on different carbon sources.

